# DNA Sequence Trace Reconstruction Using Deep Learning

**DOI:** 10.1101/2025.08.05.668822

**Authors:** Ben Cao, Lei Xie, Zhiqiang Liu, Xue Li, Bin Wang, Shihua Zhou, Pan Zheng, Qiang Zhang

**Affiliations:** The Key Laboratory of Advanced Design and Intelligent Computing, Ministry of Education, School of Software Engineering, Dalian University, Dalian 116622, China; School of Computer Science and Technology, Dalian University of Technology, Dalian 116024, China; Department of Accounting and Information Systems, University of Canterbury, Christchurch 8140, New Zealand

## Abstract

Deciphering DNA sequences is fundamental to unlocking the mysteries of life, but the high dimensionality and complexity of biological sequence data significantly hinder knowledge discovery. In particular, the challenges of sequence length, repetitive regions, and structural complexity make it difficult to directly reconstruct complete DNA sequences from raw data. Therefore, this paper proposes a DNA sequence trace reconstruction model, DNARetrace, which performs preprocessing and dataset construction, and then employs a Bidirectional Fourier-Kolmogorov-Arnold Network (Bi-FKGAT), using an extremely unbalanced loss function for link prediction, so as to reconstruct the original DNA sequence. In multi-angle experiments using both simulated and real data, DNARetrace successfully reconstructs DNA sequence traces across large-scale datasets derived from various DNA sequencing methods, overcoming the bias of current approaches toward specific sequencing platforms, and achieves competitive outcomes in DNA storage and genomics downstream tasks. We further validated the expandability of the proposed methods in DNA sequence classification and metagenomic binning tasks. In summary, DNARetrace is compatible with various sequencing scenarios; it reduces the difficulty of discovering novelty knowledge directly from high-complexity raw data, and it provides a reusable tool to accelerate DNA sequence processing and applications.

## Introduction

DNA sequences are the main bearers of genetic material, and their study and analysis are significant in disease causation, gene function analysis, and personalized medicine [1, 2]. In recent years, with the recent development of high-throughput sequencing technology, it is possible to obtain a large amount of DNA sequence data with unprecedented speed and precision [3–6], including long-read high-error nanopore sequencing, short-read low-error NGS, and high-quality and high-cost PacBio HiFi (HIFI) [7–9]. With the support of HIFI sequencing, the complete assembly of the human genome from telomere to telomere has been achieved for the first time [10]. However, long reads and accurate HiFi sequencing technology cannot read the complete genome at one time, and the original DNA sequence still must be restored by splicing the genome fragments [11, 12]. Moreover, the most used NGS cannot generate a single copy of each DNA sequence during sequencing, but it can only generate an indeterminate number of error-containing multi-copies, making it difficult to directly reconstruct the original DNA sequence [13]. Therefore, direct knowledge discovery from raw data has become a challenge for the efficient use of biological sequences, and because of different lengths of reads and error rates in sequencing technology, the current solution is weak in terms of scalability [14, 15].

Clustering and partitioning for DNA sequencing data can reduce inter-sequence interference and are indispensable as preprocessing for sequence reconstruction [16, 17]. However, the ability of current DNA clustering and partitioning methods to reconstruct the original sequences is closely related to the clusters formed by the corresponding noisy copies and thus is highly affected by DNA rearrangements and breaks. Hence, achieving perfect clustering or partitioning is challenging [17, 18]. Also, large-scale DNA sequencing data can have tens of millions, or even hundreds of millions, of nodes, in which case even the best-performing assembly methods cannot avoid high complexity in assembly graph construction [19]. Deep learning has advantages when dealing with complex data, but the highly repetitive and error-prone nature of DNA sequences limits performance. Although some methods for genome assembly have been based on graph neural networks (GNNs) [20, 21], these tend to focus on the statistical characteristics of DNA fragments and ignore the relationships between them. Therefore, they are only applicable to high-quality data obtained by HIFI sequencing but not to the highly erroneous and complex data obtained by Oxford Nanopore Technologies (ONT) sequencing, which cannot effectively ensure the integrity and accuracy of DNA sequences. As a result, the field lacks direct preference-free processing of raw data from various sequencing platforms.

A DNA sequence can be treated as a string including special information, and subsequences are rearranged and combined by trace reconstruction [22] to achieve the reconstruction and reduction of the original sequences. This paper proposes DNARetrace, a DNA sequence trace reconstruction model that does not require complex preprocessing operations such as clustering or partitioning and that can directly use raw data to achieve high-quality reconstruction of the original DNA sequence. DNARetrace performs preprocessing and dataset construction (Fig. 1A) and then employs a Bidirectional Fourier-Kolmogorov-Arnold Network (Bi-FKGAT) for link prediction to reconstruct the original DNA sequence (Fig. 1B). DNARetrace achieves the automatic conversion of wet lab data into graph structure data (Fig. 1D) by integrating multi-platform sequence alignment tools, diverse DNA fragment graph generation, and efficient labeling of DNA fragment graphs. This overcomes the problems of data scarcity and distribution bias from the root by integrating multi-source wet lab data instead of simulation data, thus ensuring the data credibility of model learning.

**Figure 1.**
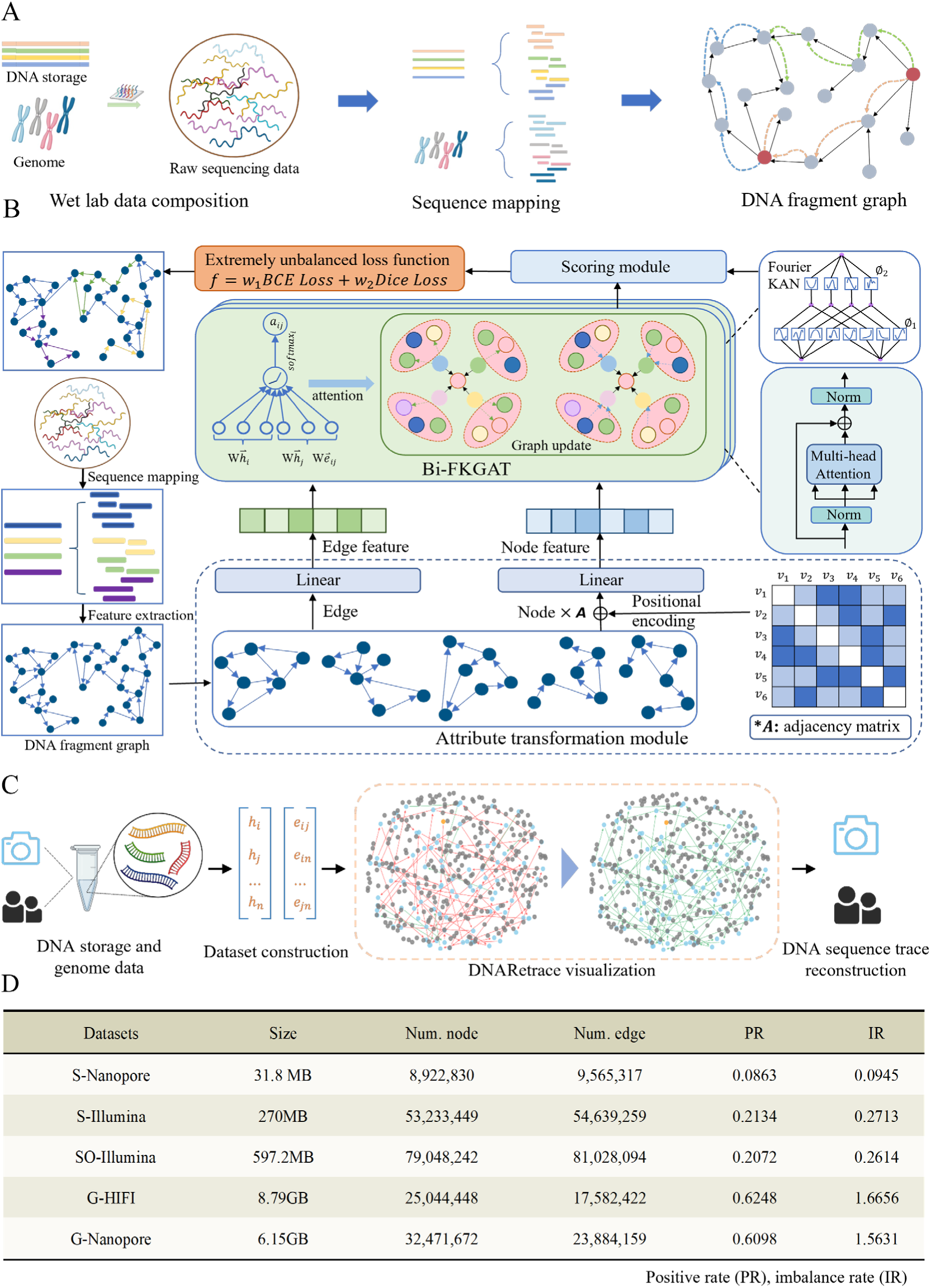
Overview of DNARetrace. (A) Dataset construction, converting wet lab data into graph format suitable for deep learning; (B) RetraceProcessor framework, consisting of an attribute conversion module, a scoring module, and a Bi-FKGAT layer, is used to learn the topological features of DNA fragment graph to achieve efficient link prediction; (C) DNARetrace main structure, application scenarios (DNA storage data reconstruction and genome assembly), and interpretability analysis; (D) Raw sequencing datasets constructed and used in this study. S-Nanopore: DNA storage dataset; 10,000 synthetic sequences (110 nt), sequenced with ONT MinION. S-Illumina: DNA storage dataset; 607,150 synthetic sequences (150 nt, 20-nt primers), sequenced on Illumina NextSeq. SO-Illumina: Aged DNA storage dataset; 210,000 sequences (200 nt, 18-nt primers), subjected to 70 °C incubation for 56 days, sequenced on Illumina HiSeq. G-HIFI: Human genome dataset; PacBio CCS data of CHM13 cell line (SRX5633451). G-Nanopore: Human genome dataset; ONT data from T2T Consortium (SRX19306105), with rebase calling for enhanced accuracy.

DNARetrace adopts a three-stage cascaded architecture. The attribute transformation module normalizes node and edge attributes into consistent embeddings. Based on a Fourier-Kolmogorov-Arnold network (FKAN), Bi-FKGAT captures complex connectivity in DNA fragment graphs. The scoring module decodes the fused features from Bi-FKGAT to filter out interference edges, enabling accurate link prediction. Bi-FKGAT addresses the unidirectional neighborhood aggregation defect of GNN-based studies [20, 21] and aggregates node input and output edge information through a bidirectional attention mechanism to enhance local structure modeling. FKAN is used to build a deep network to overcome over-smoothing [23], extracting the long-range topological features of the DNA fragment graph. To address the extreme data imbalance issue in DNA sequence trace reconstruction, DNARetrace improves the accuracy of link prediction through the adoption of a loss function combining binary cross-entropy (BCE) loss and dice-focal (Dice) loss. In addition, DNARetrace is interpretable to a certain extent. Fig. 1C shows the trace reconstruction process of various DNA sequences, which enhances the intuitive understanding of the reconstruction process and results. The details of all the datasets constructed in this paper are presented in Fig. 1D. Supplementary Section 3-6 demonstrates that the data are unbiased and illustrates the impact of errors on graph construction.

In experiments, DNARetrace achieved a 96.55% sequence recovery rate and a 98.89% base recovery rate in DNA storage, with an F1 score and MCC more than 10% greater than those of state-of-the-art models. For genome tasks, DNARetrace achieved the highest F1 score and MCC (97% and 92.28%, respectively). In genome assembly applications, it produced the most contiguous assembly, with an NA50 of 3.21 Mbp, while reducing the mismatch rate by 40.79%, demonstrating superior trace reconstruction performance. Overall, DNARetrace provides an efficient, accurate, reusable, and highly interpretable solution for knowledge discovery in DNA sequences.

## Results and Analysis

### Overview of Results

The study of DNA sequences has far-reaching impacts and wide applications in many fields, but the high dimensionality and complexity of biological sequence data, especially base errors and DNA assembly complexity, significantly hinder development [24]. To address these issues, we propose DNARetrace, an efficient and interpretable DNA sequence trace reconstruction model. By leveraging graph-based representations and the graph neural network Bi-FKGAT, DNARetrace effectively captures complex relationships within sequencing data, enabling high-precision data reconstruction and continuous assembly.

We carried out experiments with two types of DNA sequence representative tasks, synthetic DNA sequences (DNA storage) (Fig. 2) and natural DNA sequences (genomes) (Fig. 3), and we confirmed the diversity of DNARetrace by applying it to DNA sequence classification and metagenomic binning tasks. DNARetrace consistently outperformed state-of-the-art models across various tasks in our experiments, showcasing its performance and broad applicability.

**Figure 2.**
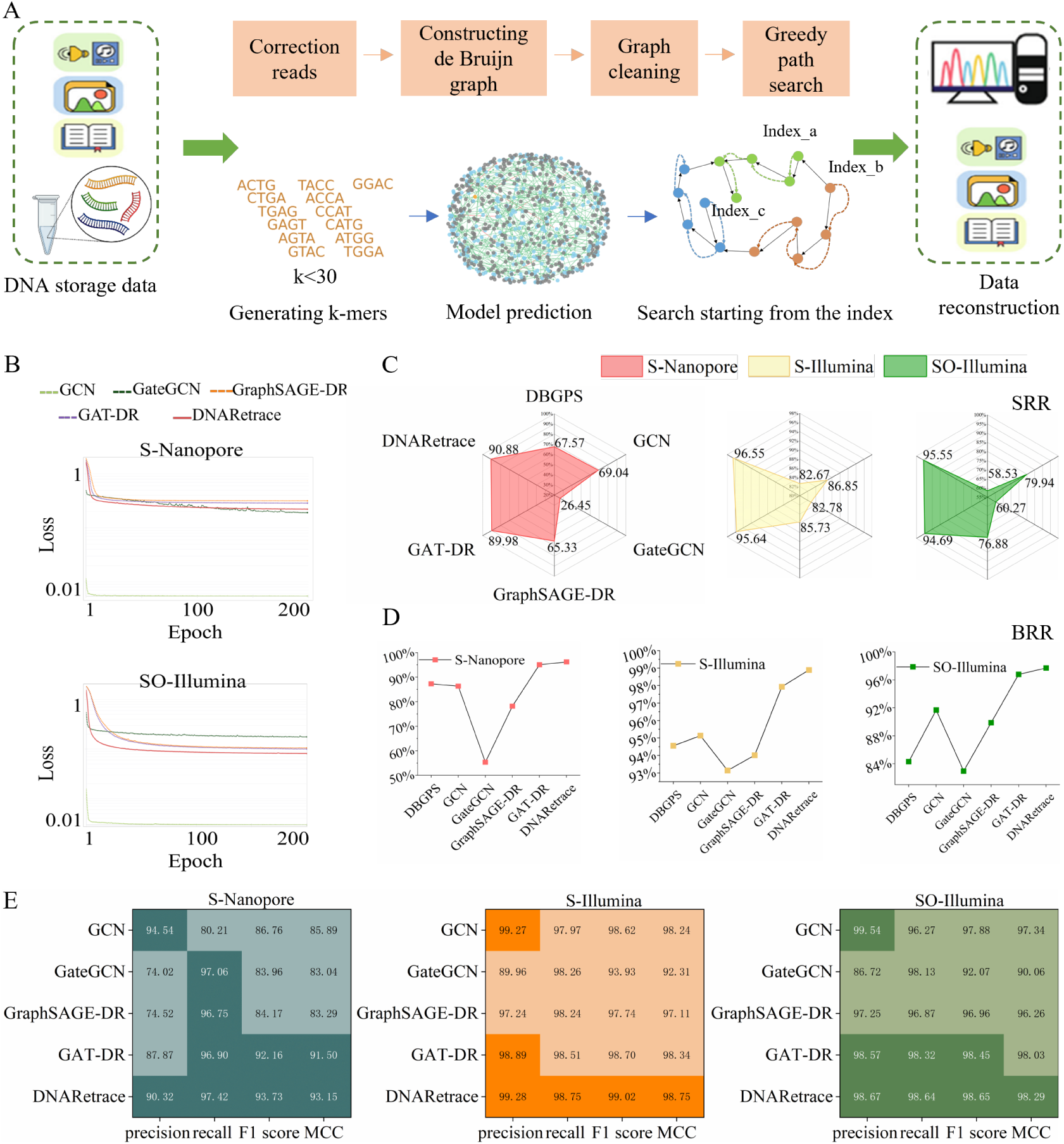
Process and performance evaluation of DNA storage data reconstruction. (A) Reconstruction process for DNA storage data: heuristic methods (top), DNARetrace (bottom), Index_a, Index_b, and Index_c indicate the starting positions for DNA sequence assembly, derived from the encoding-stage design; (B) Evolution of loss function during training; (C) Comparison of sequence recovery rate (SRR) with representative method; (D) Comparison of base recovery rate (BRR) with representative method; (E) Performance comparison of models on DNA storage datasets based on precision, recall, F1 score and MCC.

**Figure 3.**
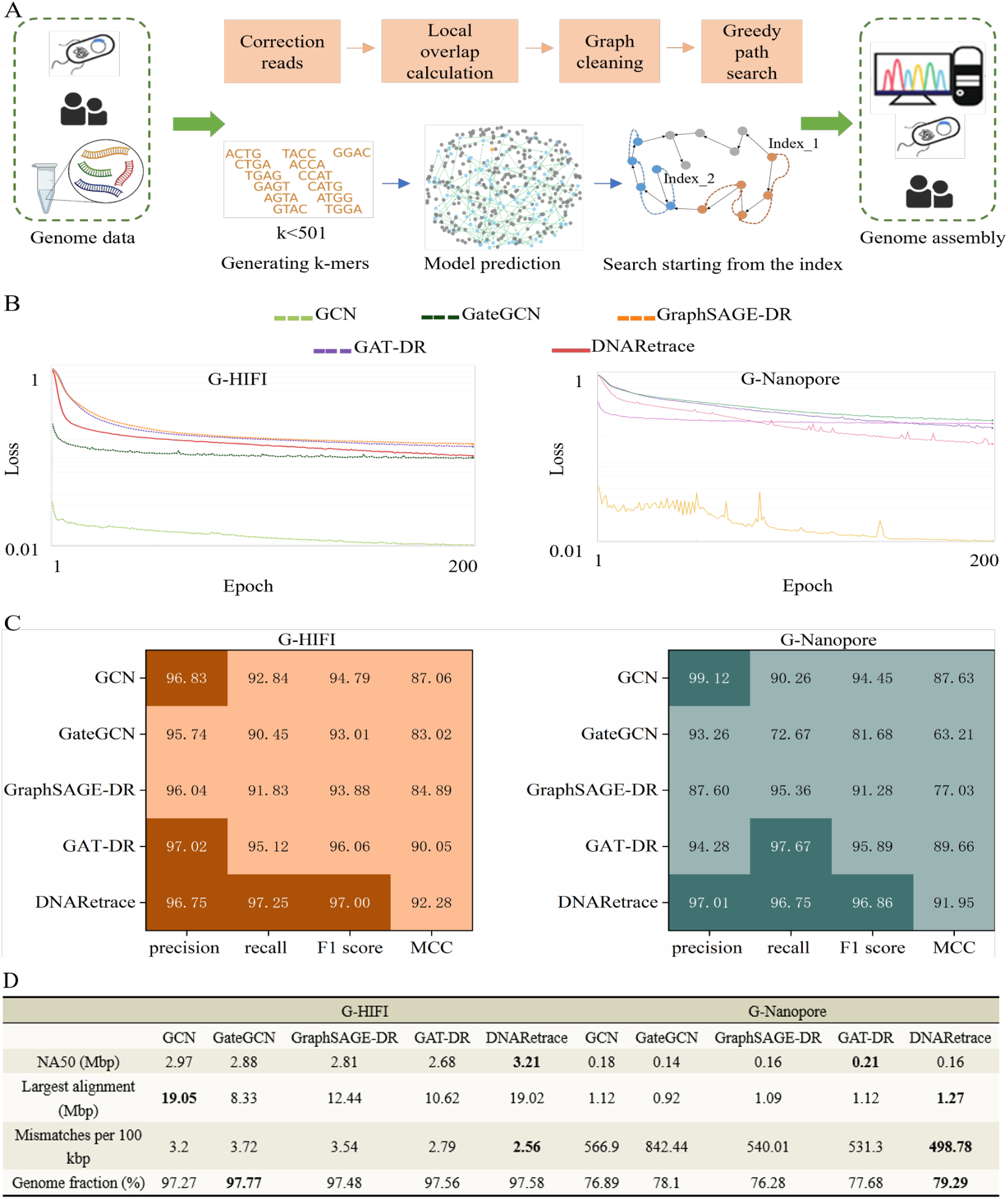
Process and performance evaluation of Genome assembly. (A) Comparison between genome assembly process of DNARetrace (bottom) for genome data and that of heuristic methods (top). Index_1 and Index_2 represent the starting points of genome assembly, which are nodes with zero in-degree; (B) Evolution of loss function during training; (C) Performance comparisons of models on genome datasets; (D) Comparison of assembly results on the human genome dataset, based on commonly used assembly evaluation metrics.

### DNARetrace for DNA Storage Data Reconstruction

The DNA molecule is widely considered to be an ideal data storage medium due to its excellent stability and high storage density [25]. However, due to the inherent limitations of synthesis and sequencing technologies and the need for long-term data preservation, there is a risk of data corruption. This highlights the necessity of reconstructing DNA storage data, which is a core application scenario of DNARetrace.

The DNA sequences carrying data in DNA storage are artificially synthesized and are selected as one of the tasks of DNA sequence reconstruction in this paper. Artificially synthesized DNA sequences have the characteristics of short length and few unwanted patterns [26]. Figure 2A shows the process of DNA storage data reconstruction using the traditional GateGCN-based method [20, 21, 27–29] and the proposed DNARetrace. Traditional methods are similar to a deep search, using the index set by the original DNA sequence as the starting point and the length of the original DNA sequence as the termination condition (Fig. 2A). The DNARetrance reconstructs the original DNA sequence based on graph learning. In this process, k-mer is used to divide the sequence into substrings containing k bases. These k-mers serve as the basic building blocks of nodes or edges in the graph, expressing the connection relationship between sequences in the graph structure. The selection of parameter k directly affects the topological structure of the graph and the reconstruction effect, which is further discussed in the Supplementary Section 1. In addition, we compare the performance of DNARetrace and other methods [20, 21, 27–29] during the training process, as well as their DNA sequence reconstruction.

### Sequence Reconstruction in DNA Storage

Since digital files are usually represented as multiple DNA sequences in DNA storage, to evaluate the overall reconstruction effectiveness of the data, we use the sequence recovery rate (SRR) [30] as a criterion, i.e.,

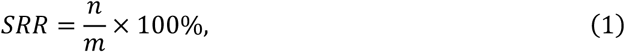

where 𝑚𝑚 and 𝑛𝑛 are the respective numbers of original and successfully recovered sequences. As can be seen from Fig. 2, the practical application results of the model are closely related to its performance. DNARetrace achieved the highest SRR on several datasets, each time exceeding 90%. In particular, on the S-Nanopore dataset, it outperformed GateGCN by more than 60%, GCN by about 20%, and DBGPS by about 23% (Fig. 2D). For DBGPS, the proposed method outperformed GateGCN and GraphSAGE on the S-Nanopore dataset, which has a high error rate, further verifying the importance of rich feature expression and neighborhood information for GNNs in the analysis of the connections between DNA fragments. This advantage of DBGPS on the S-Illumina and SO-Illumina datasets no longer exists. Although these two datasets have lower error rates than S-Nanopore, their scale is much larger, posing a path entanglement problem that seriously threatens the performance of DBGPS. SRR is used to assess the reconstruction of DNA storage data from a global perspective, and the base recovery rate (BRR) [30] is used to measure the recovery of specific sequences, where

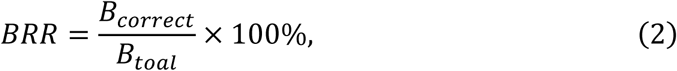

where *𝐵_𝑐orrect_* is the number of correctly reconstructed bases, and *𝐵_total_* is the total number of bases in the original sequence. As shown in Fig. 2C, BRR and SRR of the models follow the same trend, where BRR is greater than SRR, which indicates that the models recover better at the base level than at the sequence level. DBGPS exhibits a unique base recovery capability, with a slightly lower SRR than the GNN-based methods on several datasets but a significantly better BRR. For example, DBGPS outperformed GCN on the S-Nanopore dataset. DBGPS outperformed GateGCN on the S-Illumina and SO-Illumina datasets in terms of SRR, and its BRR was significantly better than that of GNN-based methods. This is mainly because DBGPS adopts greedy search in path selection, which is beneficial to base recovery, but it can easily fall into local optimization, and it has difficulty taking into account the complete recovery of the global sequence. Looking at Figs. 2C and D, DNARetrace shows the best performance at both the base and sequence levels. In particular, BRR is greater than 96% on average, which is 41% ahead of similar methods, and its excellent data reconstruction capability is seen in its robustness and the ability to reconstruct with data at an extremely large scale (Supplementary Section 8). Sequencing depth is a key variable in DNA storage data reconstruction. Hence, we also analyze the influence of sequencing depth on reconstruction efficiency (Supplementary Section 2).

### Performance evaluation of DNARetrace for DNA storage data reconstruction

We compared DNARetrace with baseline models including GCN, GateGCN, and DBGPS across multiple datasets and analyzed the results of GraphSAGE-DR and GAT-DR. As shown in Fig. 2E, DNARetrace outperforms the baseline models on all datasets. Specifically, for the S-Illumina dataset, DNARetrace performs best, with 99.28% precision (P), 98.75% recall (R), 99.02% F1 score (F1), and 98.75% MCC (M). We choose MCC instead of accuracy as the evaluation metric based on the characteristics of the dataset. As can be seen from Fig. 1D, all datasets show imbalanced characteristics. At this point, accuracy can no longer be regarded as a reliable metric, while MCC overcomes the problem of sample imbalance [31]. It can also be seen from Fig. 2E that all models perform best on the S-Illumina dataset, followed by SO-Illumina, and worst on S-Nanopore. This is mainly attributed to differences in data error rates: the lowest PR and IR were found on S-Nanopore, followed by SO-Illumina, and the highest on S-Illumina (Fig. 2E). The difference between SO-Illumina and S-Illumina stems from the fact that SO-Illumina is a product of accelerated aging experiments, which were used to validate the performance of the models in terms of their resistance to DNA degradation. The results show that DNARetrace performs well in this scenario, and MCC leads GateGCN by more than 8%.

Among the baseline models, GateGCN has the worst performance. On the S-Nanopore dataset, both the F1 score and MCC of DNARetrace are more than 10% ahead of GateGCN (Fig. 2E). The analysis shows that this gap mainly stems from the focus of GateGCN only on the edge features in the DNA fragmentation graph, while GCN also takes into account the node degree information; hence, GCN outperforms GateGCN, but not DNARetrace. From this phenomenon, we see that feature enrichment facilitates the model’s learning of graph structures. The ability of DNARetrace to surpass baseline models on multiple datasets is closely related to the feature-rich DNA fragmentation graphs constructed specifically for it by preprocessing and dataset construction. We analyze the time and memory cost in Supplementary Section 7.

We also analyzed the performance of GraphSAGE-DR and GAT-DR at processing DNA storage data. GraphSAGE-DR addresses the high computational complexity of GAT on large-scale DNA fragmentation graphs by randomly sampling neighboring nodes before polymerization. However, as shown in Fig. 2E, the performance of GraphSAGE-DR is significantly less than that of DNARetrace, and it even lags behind GCN. This is mainly because GraphSAGE employs random sampling of neighboring nodes, which results in the loss of key structural information and the failure to fully capture the complexity of DNA fragment graphs. GAT-DR shows the second-best performance in Fig. 2E, but this gradually pulls away from DNARetrace as the data error rate increases. This indicates that GAT has difficulty with the complex relationships between nodes in the presence of high data error rates. The bidirectional graph attention layer of DNARetrace can capture information of both incoming and outgoing edges, while positional encoding introduces node relative positional relationships, enabling more comprehensive modeling of node interactions.

We further visualized the loss trajectories of different models during training on the DNA storage datasets, so as to evaluate their robustness. Given the fundamental differences in the formulation and numerical scales of various loss functions, the direct comparison of their minimum loss values does not constitute an objective assessment of their effectiveness. Therefore, this study analyzes the stability and convergence performance of models by focusing on the descent trend, convergence speed, and fluctuation behavior of the loss curves.

In Fig. 2B, DNARetrace demonstrates a smooth and consistently decreasing loss curve throughout the training process, indicating a stable optimization trajectory. Although it employs the same loss function as GraphSAGE-DR and GAT-DR, DNARetrace achieves effective convergence in fewer training epochs. DNARetrace exhibits minimal variation across the entire training process, reflecting strong training stability, while GCN exhibits noticeable oscillations during the early training phase on the S-Illumina dataset, and GateGCN shows substantial fluctuations in the mid-to-late training stages on the S-Nanopore dataset.

### DNARetrace for Genome

With continuous breakthroughs in DNA sequencing technology, it has gradually become possible to obtain long and nearly perfect reads [32]. For example, the Human Genome Project was made possible by high-precision HIFI sequencing. However, the DNA fragments produced by sequencing are highly intermixed, and a precise and efficient method is required to assemble them into a complete genome, which is another important application scenario for DNARetrace. We validated the above models [20, 21, 27–29] in a practical application of genome assembly (Fig. 3).

### Sequence Reconstruction in Genome

The DNA fragment graph is input to DNARetrace; unlike the encoding stage in DNA storage where indexing can be set, the genome sequence reconstruction cannot directly select the ideal starting node. Therefore, we select the node that has a zero entry degree as the starting point, adopt a method similar to depth-first search, and extract the longest path (Fig. 3A). Due to resource constraints, we only use 4000 nodes as starting nodes for the experiment. For the final output sequences of the model, we use QUAST-LG version 5.3.0 and benchmark with T2TGenome [33] as the reference genome.

As shown in Fig. 3D, on the G-HIFI dataset, DNARetrace has a slightly shorter longest alignment length than GCN (19.02 Mbp versus 19.05 Mbp) and a slightly smaller Genome fraction than GateGCN (97.58% versus 97.77%), but these differences are not significant. However, DNARetrace generated the most contiguous assembly (NA50 = 3.21 Mbp) and achieved the lowest mismatch rate (only 2.56 mismatches per 100,000 base pairs). Hence, DNARetrace has a significant advantage in assembly precision. The G-Nanopore dataset has a higher error rate than G-HiFi, and hence, each model suffers significant performance degradation (Fig. 3D). However, benefiting from the rich features provided by preprocessing and dataset construction, DNARetrace, with its powerful bidirectional feature extraction and remote information capture, still performs well, with a 40.79% reduction in the mismatch rate compared to GateGCN. In addition, its longest alignment length and genome fraction exceed those of the baseline model. GraphSAGE-DR and GAT-DR both show some advantages in practical applications, but there is still a gap compared with DNARetrace. This also indicates the stronger capabilities of DNARetrace in feature extraction, information transfer, and model accuracy, further enhancing its performance at genome assembly tasks.

### Performance Evaluation of DNARetrace in Genome

We compare DNARetrace with baseline models GCN and GateGCN on the genome wet lab dataset and analyze the performance of GraphSAGE-DR and GAT-DR. DNARetrace performs well on the G-HIFI and G-Nanopore datasets, especially in the key metrics of F1 score and MCC, where it significantly outperforms GCN and GateGCN (Fig. 3C). On the G-HIFI dataset, the F1 score of DNARetrace reaches 97.00%, which is 3.99% ahead of the baseline model, and MCC reaches 92.28%, which is 9.26% higher than the baseline model. On the G-Nanopore dataset, the F1 score of DNARetrace is 2.41% and 15.18% higher than those of GCN and GateGCN, respectively, and the corresponding values of MCC are 4.32% and 28.74% higher than those of GCN and GateGCN, respectively. DNARetrace has above advantages, which is closely related to its model depth and loss function. The number of GNN layers reaches six, which can effectively capture complex features and structures, thus improving the model’s expressiveness and prediction performance. In addition, DNARetrace adopts a joint BCE and Dice loss function and improves the accuracy and stability of the model with imbalanced data by adjusting the weights and focusing on difficult-to-classify samples.

In Fig. 3C, GAT-DR shows competitive performance due to the attention mechanism, which effectively differentiates the attention level of neighboring nodes, and it can more effectively capture complex relationships and features. GraphSAGE-DR performs more poorly than GAT-DR due to sample learning, but it still outperforms GateGCN, which utilizes only edge features. In the baseline model, GCN usually outperforms GateGCN, as GCN can more comprehensively integrate node features and edge information, and thus exhibits stronger expressive and predictive abilities than GateGCN.

Fig. 3B shows the changes in training loss of each model on the human genome dataset. Since G-Nanopore has a significantly greater error rate than G-HIFI, the training stability of each model on this dataset generally decreases, especially for GCN, where the loss curve fluctuates violently. In both datasets, DNARetrace always shows a lower loss value than GraphSAGE-DR and GAT-DR, indicating stronger feature extraction capabilities. For G-HIFI, the loss decrease process of DNARetrace is smoother than that of GateGCN; on G-Nanopore, although DNARetrace is slightly less stable than GateGCN, its loss always maintains a downward trend. However, GateGCN experiences a loss rebound in the later stage of training, showing a certain degree of overfitting, further demonstrating the robustness and generalization ability of DNARetrace in complex sequencing environments.

### Expandability Analysis of Bi-FKGAT Within DNARetrace

A graph, as a complex data structure, can effectively capture local and global non-adjacent information in data. With the wide application of graph structures in bioinformatics, they have been explored in recent years for tasks such as DNA sequence classification [34] and metagenomic binning [35]. Within DNARetrace, although designed for link prediction tasks for DNA sequence trace reconstruction, Bi-FKGAT, as a kind of GNN, has a powerful ability to learn graph structures, which makes it suitable for classification tasks. We model DNA sequence classification and edge classification tasks, so as to compare Bi-FKGAT with the baseline models.

In the DNA sequence classification task, the composition is mainly realized through the relationship between subsequences [34]. First, the sequence is decomposed into a set of subsequences, which are represented as nodes, between which edges are created. Different sequences can be connected to each other through common subsequences. Finally, semi-supervised learning is applied on this graph and a cross-entropy loss function is used to train the model. Fig. 4A shows the results of the model on the Anticancer peptides dataset [34]. It can be seen that Bi-FKGAT outperforms all baseline models, with the highest F1 score of 91.37%, exceeding that of the second-best DNA-GCN [36] by 7.55%. The success of Bi-FKGAT lies in its ability to capture higher-order, complex relationships in graph structures. In addition, it utilizes a multi-head attention mechanism to enhance sequence representation learning, making it superior to DNA-GCN, which shows competitive performance due to its heterogeneous graph-based structure, allowing information passage between sequences and subsequences. Also, Fig. 4A shows that support vector machine (SVM) outperforms deep learning models such as the recurrent convolutional neural network (RCNN) and bidirectional long short-term memory (BiLSTM), perhaps because deep learning models require more data and use relatively small datasets, thus limiting their performance.

**Figure 4.**
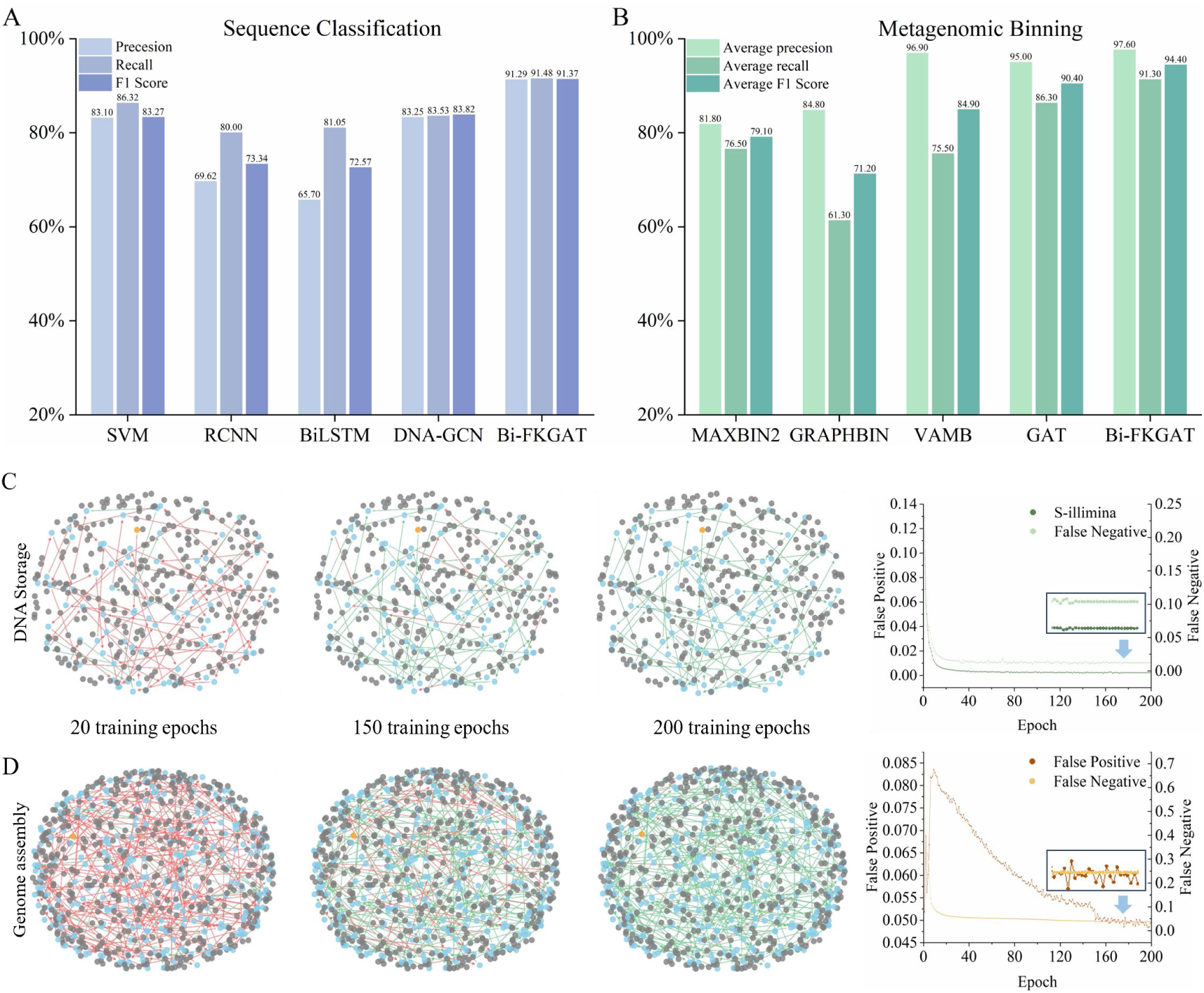
Expandability and interpretability analysis. (A) Bi-FKGAT was used to perform sequence classification tasks and comparison with baseline models using Anticancer peptides dataset, which contains 949 one-letter amino-acid sequences representing peptides and their four anti-cancer activities; (B) Bi-FKGAT was used to perform the task of metagenomic binning and comparison with baseline models, using Strong 100 dataset, which contains reads from 100 strains, corresponding to 50 species, with read abundances randomly generated; contigs were generated using metaflye [37]; (C) Visualization of DNA sequence trace reconstruction process on S-Illumina dataset across different training epochs, along with corresponding changes in false-positives and false-negatives; (D) Visualization of DNA sequence trace reconstruction process on G-HIFI dataset across different training epochs, along with corresponding changes in false-positives and false-negatives. (start nodes: yellow; internal nodes of paths: blue; noise nodes: gray; paths that failed to be predicted: red edges; paths that were successfully predicted: green)

Bi-FKGAT can also be applied to unsupervised tasks, with metagenomic binning being a prime example. Learning contig representations via assembly graphs, this task is well suited to implementation using GNNs. Lamurias et al. [35] proposed a framework to utilize assembly graphs for metagenomic binning. In this framework, we use the Strong 100 dataset [35] to compare Bi-FKGAT with current mainstream binning methods [35] such as MAXBIN2, VAMB, and GRAPHBIN and with GAT [29], so as to demonstrate its advantages over similar models. Fig. 4B shows that GNN outperforms other methods in almost all metrics. This is because most methods rely on nucleotide composition, abundance features, or local contig features, treating sequences as independent data points without considering that some may originate from contiguous genomic regions. In contrast, Bi-FKGAT obtains neighborhood-aware embeddings for each contig by training a graph neural network on the assembly graph to more accurately distinguish reads from different strains. In addition, Bi-FKGAT can scale up to six layers, and it learns bidirectionally from neighboring nodes to capture more global information. Bi-FKGAT outperforms GAT by virtue of multi-layer information dissemination and bidirectional learning of neighboring nodes, and it ultimately performs best, with average precision (AP) of 97.6%, average recall (AR) of 91.3%, and an F1 score of 94.4%.

### Interpretability analysis of DNARetrace

The weak visualization of graph data compared to images and text adds to the difficulty of interpreting the process performed by DNARetrace in DNA sequence trace reconstruction. To address this, we carry out preprocessing and dataset construction to generate the datasets S-Illumina (Fig. 4C) and G-HIFI (Fig. 4D) and demonstrate the process of link prediction by visualizing DNARetrace. Specifically, we continuously track and display the path changes of one of the sequences in the DNA fragmentation graphs constructed from the two datasets and observe its dynamics under different training cycles (start nodes: yellow; internal nodes of paths: blue; noise nodes: gray; paths that failed to be predicted: red edges; paths that were successfully predicted: green edges).

The length of the original DNA sequence of S-Illumina is 150 bp, which is much less than that of G-HIFI, and thus, the number of nodes in the path is relatively small, which is why Fig. 4C is sparser than Fig. 4D. As shown in Fig. 4, the number of correctly predicted edges gradually increases with the training period. However, DNARetrace has a higher percentage of incorrectly predicted edges on G-HIFI when the training period reaches 150 epochs. From our analysis, this is mainly because the DNA fragmentation graph generated from G-HIFI is sparse, which leads to slower convergence. In addition, DNARetrace still has incorrectly predicted edges for G-HIFI when the training period reaches 200 epochs. This is due to the extremely long target path length of genome assembly, which increases the difficulty of model prediction.

We also observed the changes in false-positives and false-negatives of the model at each stage. Fig. 4C and Fig. 4D show that, with the increase of training cycles, false-positives and false-negatives have a downward trend, which is consistent with the expectation that the model performance gradually converges and optimizes. The difference is that on G-HIFI, the model has a significant increase in false-positives in the early stage of training. This is because the number of positive samples in the dataset is relatively large, which makes the model overly sensitive to the positive class in the early training stage and tends to judge more samples as positive, thus causing a surge in false-positives. As the training progresses, the model gradually learns more effective discrimination boundaries, and the false-positive rate decreases accordingly.

### Ablation Study

To gain deeper insights into the contributions of individual components within DNARetrace, we conducted ablation studies on the S-Illumina and G-HIFI datasets, chosen for their low error rates in DNA storage and human genome contexts. Specifically, we investigated the impact of varying the number of bidirectional graph attention (GAT) layers, the inclusion of the FKAN module, and the bidirectional attention mechanism itself. The summarized results in Table 1 reveal several key findings. Firstly, reducing the number of bidirectional GAT layers within the Bi-FKAN module to just two led to measurable decreases in Matthews Correlation Coefficient (MCC) of 0.15% on S-Illumina and a more substantial 3.14% on G-HIFI. Secondly, the removal of the FKAN module (w/o FKAN) consistently resulted in reduced performance across both F1 score and MCC metrics. Most significantly, eliminating the bidirectional attention mechanism (w/o Bi) caused a pronounced drop in both F1 score and MCC, exceeding the impact observed from removing FKAN. This underscores the bidirectional mechanism’s critical role in effectively learning the connectivity between DNA fragments. Furthermore, Table 1 highlights that performance degradation across all ablation conditions was consistently more severe on the G-HIFI dataset compared to S-Illumina. We attribute this heightened sensitivity to the greater sparsity inherent in the G-HIFI data (Fig. 1D), which necessitates the deeper representational capabilities provided by multiple GAT layers, the FKAN module, and the bidirectional attention mechanism to adequately capture complex DNA fragment features. Comparative analyses against common GNN architectures, detailed in Supplementary Section 9, further substantiate the unique advantages offered by the full Bi-FKGAT design.

**Table 1:** Effectiveness of bidirectional graph attention layers (𝐿𝐿), FKAN, and bidirectional attention mechanism.

| Model | S-Illumina |  |  |  | G-HIFI |  |  |  |
| --- | --- | --- | --- | --- | --- | --- | --- | --- |
|  | P | R | F1 | M | P | R | F1 | M |
| RetraceProcessor ($L=2$) | 99.14 | 98.67 | 98.90 | 98.60 | 96.51 | 94.91 | 95.70 | 89.14 |
| RetraceProcessor ($L=4$) | 99.22 | 98.71 | 98.96 | 98.68 | 96.70 | 96.63 | 96.66 | 91.45 |
| RetraceProcessor (w/o FKAN) | 99.20 | 98.68 | 98.94 | 98.65 | 96.51 | 96.64 | 96.57 | 91.18 |
| RetraceProcessor (w/o Bi) | 99.16 | 98.67 | 98.92 | 98.63 | 96.36 | 96.50 | 96.43 | 90.81 |
| RetraceProcessor | **99.28** | **98.75** | **99.02** | **98.75** | **96.75** | **97.25** | **97.00** | **92.28** |

## Materials and Methods

### Experimental design

#### Datasets

We evaluated DNARetrace on five wet lab datasets (as shown in Table 3), which originate from DNA storage and human genome domains, and demonstrated their significant heterogeneity in both scale and imbalance characteristics.

1. **DNA storage**.

1. S-Nanopore. The sequencing file was provided by Srinivasavaradhan et al. [38], and the data were encoded into 10,000 DNA sequences of length 110 nucleotides for storage. The DNA sequence was synthesized by Twist Bioscience, amplified via polymerase chain reaction (PCR), ligated to ONT adapters, and sequenced on the MinION platform.
2. S-Illumina. The sequencing file was provided by Orgick et al. [39], who mapped the digital files into 607,150 DNA sequences of 150 nucleotides in length during the encoding phase, with 20 nucleotides primer fragments at both ends. The DNA sequence was synthesized by Twist Bioscience, processed using the Illumina TruSeq Nano ligation protocol, and sequenced on the Illumina NextSeq platform.
3. SO-Illumina. The sequencing file was provided by Song et al. [27], and the data were encoded into 210,000 DNA sequences of 200 nucleotides in length, with primer fragments of 18 nucleotides at both ends of each sequence. The DNA sequence was synthesized by Twist Bioscience, and sequencing libraries were prepared from purified PCR products using the Illumina TruSeq DNA PCR-Free Library Preparation Kit and sequenced on the Illumina HiSeq platform. It should be noted that DNA serves as a long-term storage medium, and to further validate the actual performance of the model in terms of tolerance to DNA degradation, accelerated aging of sequencing files was performed in this study. The purified PCR products were continuously incubated in an elution buffer at 70°C for 56 days.

Following preprocessing and dataset construction, we converted each dataset into 10 graphs of similar size in a 6:2:2 ratio for training, validation, and testing, respectively.

2. **Human genome**

1. G-HIFI. PacBio Circular Consensus Sequencing (CCS) of the CHM13 human haploid cell line for genome assembly and variant detection. The sequencing file is from the publicly available dataset SRX5633451.
2. G-Nanopore. The original FAST5 data were generated by the Telomere-to-Telomere (T2T) Consortium and released into the public domain under a CC0 license. The sequencing file is from the publicly available dataset SRX19306105. Notably, this dataset underwent rebase calling to improve sequencing accuracy and data quality. As shown in Table 3, the positive rate (PR) and imbalance rate (IR) of this dataset are comparable to those of G-HiFi, demonstrating its high reliability.

To ensure the reliability and accuracy of the experimental results, we selected 10 chromosomes preprocessing and dataset construction. The data of chromosomes 3, 4, 5, 6, 7, and 8 were used for model training, the validation data came from chromosomes 9 and 10, and the model was tested on chromosomes 11 and 12. There is no preference for these chromosomes, except that their data are sufficient.

#### Baselines

We compare DNARetrace with four types of baseline models: (1) the GateGCN-based method [20]; (2) the GCN-based method [21]; (3) heuristic-based methods used for DNA storage data reconstruction, such as DBGPS [27]; (4) GraphSAGE [28] and GAT [29] in the DNA sequence track reconstruction task, which we call GraphSAGE-DR and GAT-DR, respectively.

#### Reproducibility

We use six bidirectional graph attention layers, each consisting of an FKAN. For the bidirectional graph attention layer, a multi-head attention mechanism is used, and attention updates based on message passing are adopted. We consider the number of attention heads ℎ = 8, hidden feature vector ℎ = 32, and weight decay 𝑑𝑑 = 1𝑒𝑒^−5^ and use Adam to optimize DNARetrace. We train the model using an early stopping strategy until there is no improvement in the Matthews correlation coefficient (MCC) [31], as verified in 200 epochs on a single NVIDIA A40 48GB GPU.

### DNARetrace design

DNARetrace converts wet lab data into a format suitable for graph neural network training through sequence mapping, DNA fragment graph construction, and rapid annotation (Fig. 1A). It ensures data reliability by integrating multi-source wet lab data, fundamentally overcoming data scarcity and distribution bias. Bi-FKGAT is used to achieve efficient interactive feature representation aggregation, thereby realizing accurate link prediction and providing strong support for DNA sequence trace reconstruction (Fig. 1B). DNARetrace employs a joint loss function to ensure overall prediction stability while enhancing sensitivity to minority-class edges, effectively reducing the bias caused by edge imbalance, and improving the accuracy of DNA sequence trace reconstruction.

### Dataset Construction

Generating the dataset can be roughly divided into three tasks (Fig. 1A): (1) generate read clusters by mapping reads to reference sequences by **Sequence mapping**; (2) **Construct a DNA fragment graph** for each read cluster; and (3) perform the path search in the DNA fragment graph, and quickly map the edges that the path passes through to the reference sequence to complete the ground-truth pathfinding by **Constructing labeling edges**.

#### Sequence mapping

Given the importance of data quality to deep learning models, we propose a method that can directly transform wet lab data into a format suitable for training graph neural networks, thereby improving data utilization efficiency. Ideally, when performing DNA sequence trace reconstruction, each reference sequence (the original DNA sequence from the DNA storage or genome) has its corresponding independent set of reads, which are represented by multiple independent DNA fragment graphs. However, sequencing data often contain reads of multiple reference sequences, which can form a large DNA fragment graph. To ensure the consistency of the training data, we use minimap2 [40] and BWA [41] to map the reads to the corresponding reference sequences to form read clusters.

In this paper, the wet lab data are divided into two categories: (1) DNA storage, generated by nanopore and Illumina sequencing technologies, as provided by Srinivasavaradhan, Organick, and Song et al.; and (2) data from the Human Genome Project [10], where the main sequencing technologies include HiFi and ultra-long nanopores. For DNA storage data, each cluster contains 10,000 reads of the original sequence, while for genome data, each cluster corresponds to reads of a chromosome. From these read clusters, we construct DNA fragment graphs that enable the generation of ground-truth paths for supervised learning.

#### Construction of DNA fragment graph

The goal of DNA sequence trace reconstruction is to extract the links between fragments in a DNA fragment graph for reference sequence recovery. However, there are problems related to the existence of few links and difficulty in feature extraction in the current DNA fragment graph. To ensure that the model learns complete information about the data, we removed all simplification operations [42], and finally each dataset contained ten de Bruijn graphs.

#### Construction of labeling edges

There are a large number of edges in the DNA fragment graph, and direct traversal generates many sub-optimal sequences. In addition, due to synthesis, sequencing, and environmental factors, sequencing data inevitably generate errors [43], which make it difficult to cross certain regions of the DNA fragment graph and thus make it impossible to mark edges.

To solve this problem, we adopt a path search method based on randomly generating many starting points to achieve a comprehensive traversal of the DNA fragment graph, and as usual for sub-optimal labeling 0 and ground-truth labeling 1. However, this method generates many edges, which significantly increases the time cost of labeling. To accelerate this process, we introduce a k-ordered induced suffix sorting technique [44] for the efficient construction and utilization of k-ordered FM indexes, so as to efficiently perform pattern matching of edge sequences in the reference sequence. Through an accurate and rational labeling process, we construct a well-established dataset that can be used to perform downstream tasks.

### DNARetrace framework

#### Attribute transformation module

In the DNA fragment graph, the sequencing data are divided into multiple DNA fragments, which are used as nodes in the graph (Fig. 1B). The connection relationship between nodes is established during the assembly process through the overlap between fragments. In this case, directly using the base sequence of the DNA fragment itself to learn the connection relationship has little effect. Therefore, introducing statistical features becomes an efficient way to represent data. In addition to the commonly used attributes such as node abundance, edge abundance, and node degree, we introduce a new feature attribute for the task of DNA sequence trace reconstruction, the abundance similarity value. Since the nodes in the de Bruijn graph are generated by sliding windows, in theory, adjacent nodes on the real path should have similar abundance values. The abundance similarity value is based on this observation, and it helps guide the direction selection of path prediction. The de Bruijn graph constructed by DNARetrace has the following structural properties: (1) input node degree; (2) output node degree; (3) node abundance; (4) edge abundance; and (5) abundance similarity value. To more deeply capture the connectivity between DNA fragments, we designed a node and edge attribute transformation module specialized for the DNA fragment graph, which converts the node attribute 𝑋 ∈ ℝ^𝑁×𝑑^𝑣 and edge attribute 𝐸∈ ℝ^𝑁×𝑑^𝑣 into d-dimensional representations. As described in the preprocessing and dataset construction, each node has three attributes and each edge has two attributes, so set *𝑑_𝑣_* = 3 and *𝑑_e_* = 2.

In the context of link prediction, some features of a node may affect the affinity to another node (due to shared attributes or latent factors); so, we consider the attributes of neighboring nodes to complete the similarity metric and update the adjacency matrix (𝑆𝑆) to incorporate the features of neighboring nodes, as follows:

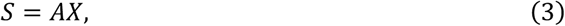

where 𝐴 ∈ ℝ^𝑁×𝑁^is the adjacency matrix. Unlike the common practice in graph neural networks, when constructing the DNA fragment graph, we avoid symmetry ambiguity by partitioning the DNA fragments into odd lengths, so the identity matrix cannot be added to the adjacency matrix.

In DNA sequences that contain data or species information, DNA fragments representing specific information (e.g., data indexes, error-correcting code redundancy, core promoters, and translation initiation sites) are usually located in fixed regions [45–48]. Hence, the nodes they represent are more influential. PageRank provides a measure of this influence, thus facilitating better link prediction. To obtain the impact of structural importance, we compute the positional encoding based on the PageRank value of each node. Formally, the page rank value of node 𝑣_𝑖_ is denoted as Pr(𝑣_𝑖_), and we compute its page rank-based position encoding (PRE) as follows:

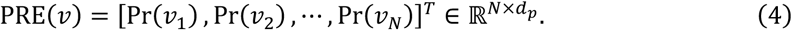

By aggregating the attributes of nodes represented by DNA fragments with positional encodings, we enrich the understanding of the DNA fragment graph by graph neural networks. Since edges are connections of nodes, we need only transform their properties into the appropriate dimension and update them according to the nodes in a specific network. The conversion formulae regarding node and edge attributes (ℎ and 𝑒) are as follows:

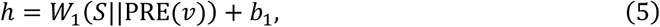

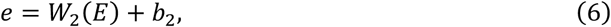

where (·||·) denotes concatenation, 𝑊_1_ and 𝑊_2_ are learnable matrices, and 𝑏_1_ and 𝑏_2_ are learnable parameters.

#### Bi-FKGAT

Graph neural networks are feasible as a powerful graph feature extraction tool for link prediction tasks. However, the number of nodes and edges of DNA fragment graphs can be in the tens of millions, and multi-layer graph neural networks are usually required to learn their complex features. Most models extract features via multi-layer perceptron (MLP) and incorporate activation functions between layers to enhance nonlinearity. However, MLP parameter sizes do not grow linearly with the number of layers, and lack interpretability [49], while activation functions (such as ReLU) may limit the characterization ability and hinder the learning of complex features [50].

To better learn the node features in DNA fragment graphs, we propose Bi-FKGAT (Fig. 1B), which introduces FKAN to reduce the use of learnable matrices and activation functions while enhancing the learning of DNA fragment graphs through a bidirectional graph attention layer that incorporates a multi-head attention mechanism. This design is capable of assigning different weights to outgoing and incoming edges, aggregating information from different directions, and hence more accurately capturing the upstream and downstream relationships of DNA fragments. FKAN overcomes the difficulty of training the spline function in the traditional Kolmogorov-Arnold network [51], which leads to difficult model convergence, and the Fourier coefficients [52] can be a good substitute for the spline function. The equation of FKAN (𝜙_𝐹_) is

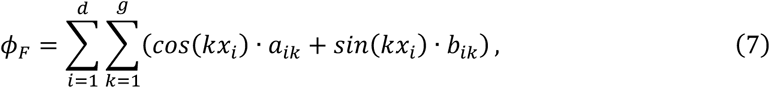

where 𝑑 is the number of dimensions of features, and *𝑎_ik_* and *b_ik_* are learnable parameters. The hyperparameter 𝑔 is the grid size, and 𝑔 = 2 is recommended.

We utilize FKAN instead of MLP as part of the feature transformation in the GAT message-passing process, i.e.,

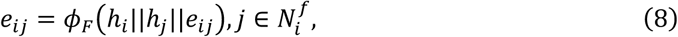

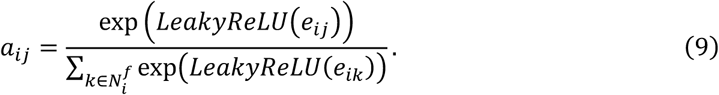

Here, the attention weights are computed based on the outgoing edges of the nodes in the forward graph, and similarly, the attention weights of the incoming edges in the reverse graph are computed as follows:

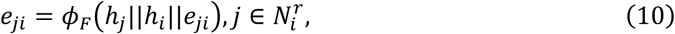

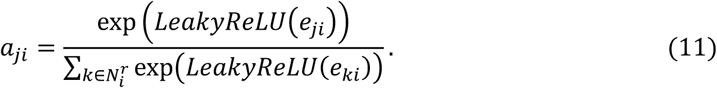

After completing the attention weight calculation, Bi-FKGAT performs a comprehensive update of the node features through the multiple attention mechanism:

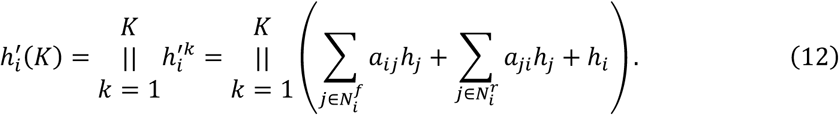

Since edges and their end nodes are closely related, edge features can be updated using node features:

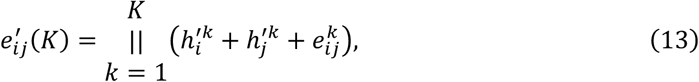

where (·||·) denotes concatenation; 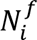 and 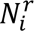 are neighboring nodes of node *i* in the forward and reverse graphs, respectively; and 𝑘 denotes the attention head.

#### Scoring module

In link prediction tasks, a common training strategy is to optimize the model by bringing the nodes connected to the target edge closer to each other, which can be reflected by the edge scores. FKAN decodes the obtained edge representation into a probability score (Fig. 1B). The probability score *𝑝_ij_* for traversing an edge 𝑠 → 𝑗 is computed from node representation of nodes 𝑠 and 𝑗, as well as edge representations of the edge 𝑠 → 𝑗, all after the final Bi-FKGAT layer 𝐿:

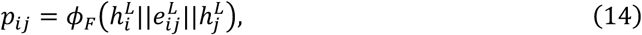

where (·||·) denotes concatenation.

### Extremely Unbalanced Loss Function Design

Although HIFI sequencing technology and high accuracy ONT sequencing technology provide high-quality genomic data, they are costly and have complex library construction processes. Even with 99% sequencing accuracy [12], there are still a large number of error edges in the DNA fragment graph, leading to an imbalance in genomic data even with high-precision sequencing technologies. Furthermore, sequencing technologies commonly used in the field of DNA storage, such as Illumina and ONT, cannot achieve the high accuracy of HIFI, which further exacerbates the imbalance between correct and incorrect edges in the graph, making the data even more imbalanced. For DNA molecules that must be stored for long periods of time, such as ancient DNA or DNA data storage, errors inevitably accumulate with long-term storage, reading, and writing, resulting in a significant increase in error edges in the DNA fragment graph, with the proportion of correct edges dropping below 10% in extreme cases. Therefore, DNA sequence trace reconstruction must specifically address the issue of extreme data imbalance (dataset characteristics are shown in Table 3).

Traditional classification loss functions have certain limitations in extremely unbalanced data environments, so we adopt a extremely unbalanced loss function to improve the robustness and accuracy of the model under such conditions,

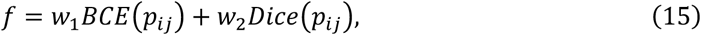

where *𝑝_ij_* is the predicted value of the model for the edge (*𝑁ode*_𝑖_ → *𝑁ode_j_*), and *BCE Loss* measures the deviation between the predicted and true values by capturing the overall distribution of the data. However, when the data is extremely unbalanced, *BCE Loss* will be affected by the unbalanced samples, and the model prediction will tend to favor the majority class. Therefore, DNARetrace introduces *Dice Loss* in the loss function, which enhances the identification of minority classes by assessing the degree of overlap between predicted and true values and effectively mitigates the negative impact of data imbalance and improves the predictive performance of the model on positive samples. By setting the weights 𝑤_1_ and 𝑤_1_, it ensures that the loss function enhances the sensitivity to the minority class with effective overall prediction performance and improves the reconstruction performance of DNARetrace in extremely unbalanced datasets.

## Conclusion

In orde to solve the problem of the high dimensionality and complexity of biological sequence data significantly hinder knowledge discovery. We introduced DNARetrace, a model designed for DNA sequence trace reconstruction. DNARetrace first conducts preprocessing and dataset construction, an then propose the Bi-FKGAT model for learning sequence features and reconstruct original DNA sequence, which can be applied to short-read DNA storage data as well as long-read genomic data, successfully realizing efficient and robust reconstruction.

The performance and robustness of DNARetrace were comprehensively evaluated in a series of comparative experiments covering DNA storage data reconstruction, genome assembly, DNA sequence classification, and metagenomic binning tasks. In DNA storage data reconstruction and genome assembly experiments, we evaluated the effectiveness of DNARetrace under complex sequencing conditions using raw sequencing data generated from Illumina, ONT, and HiFi technologies, where DNARetrace achieved a sequence recovery rate of over 96% and a base recovery rate of 98% in DNA storage data reconstruction. In genome assembly, it attained an F1 score of 97%, improved NA50 to 3.21 Mbp, and reduced the mismatch rate by 40.79% compared with the baseline model. To evaluate the generality and applicability of DNARetrace, we applied Bi-FKGAT to DNA sequence classification and metagenomic binning tasks. As show in result, Bi-FKGAT outperformed the best competing models by 7.55% and 4% in F1 score. To enhance the interpretability of the model, we conducted a visualization analysis of the link prediction process of DNARetrace and performed an ablation study to validate the roles of the bidirectional attention mechanism and FKAN in the model. DNARetrace can generate DNA fragment maps from wet experimental data, generate supervisory signals for subsequent tasks, and convert the data into a format suitable for deep learning models, while existing methods are limited by insufficient data or reliance on simulated data. Bi-FKGAT possesses powerful feature extraction capabilities, accurately analyzing DNA sequencing errors and complex DNA fragment relationships in large-scale data and efficiently integrating key topological information to optimize the DNA sequence trace reconstruction process, thereby enhancing reconstruction accuracy and overall performance.

In the future, we will explore the ability of DNARetrace to reconstruct DNA sequence trace under higher error rates and lower sequencing depths, further expanding its applicability in complex DNA sequence tasks. Expand the applicability of DNARtrace in other synthetic biology applications, such as DNA tiles, nano-delivery systems, and DNA origami.

